# Repetitive sensory stimulation potentiates and recruits sensory-evoked cortical population activity

**DOI:** 10.1101/2024.08.06.605968

**Authors:** Leena Eve Williams, Laura Küffer, Tanika Bawa, Elodie Husi, Stéphane Pagès, Anthony Holtmaat

**Author notes:** To whom correspondence should be addressed: Dr. Leena Williams, Prof. Anthony Holtmaat.

## Abstract

Sensory experience and learning are thought to be associated with plasticity of neocortical circuits. Repetitive sensory stimulation can induce long-term potentiation (LTP) of cortical excitatory synapses in anesthetized mice; however, it is unclear if these phenomena are associated with sustained changes in activity during wakefulness. Here we used time-lapse, calcium imaging of layer (L) 2/3 neurons in the primary somatosensory cortex (S1), in awake male mice, to assess the effects of a bout of rhythmic whisker stimulation (RWS) at a frequency by which rodents sample objects. We found that RWS induced a 1h-increase in whisker-evoked L2/3 neuronal activity. This was not observed for whiskers functionally connected to distant cortical columns. We also found that RWS altered whether individual neurons encoded subsequent stimulus representation by either being recruited or suppressed. Vasoactive intestinal-peptide-expressing (VIP) interneurons, which promote plasticity through disinhibition of pyramidal neurons, were found to exclusively elevate activity during RWS. These findings indicate that cortical neurons’ representation of sensory input can be modulated over hours through repetitive sensory stimulation, which may be gated by activation of disinhibitory circuits.

**SIGNIFICANCE STATEMENT:** Sensory experience and learning are thought to be associated with the plasticity of cortical synaptic circuits. Here, we tested how repeated sensory stimulation changes subsequent sensory-evoked responses, using the mouse somatosensory cortex as a model. This cortical area processes, among others, sensory information from the whiskers. We found that rhythmic whisker stimulation potentiated excitatory neuronal activity for an hour, and identified a disinhibitory interneuron-mediated mechanism that could gate this plasticity. This work increases our understanding of sensory learning and experience-dependent plasticity processes by demonstrating that cortical representations of sensory input are dynamic and are effectively modulated by repeated sensory stimulation.

## INTRODUCTION

Changes in sensory experience and perceptual learning are thought to be associated with the plasticity of cortical synaptic circuits (Feldman 2009; Chéreau et al. 2020). Sensory deprivation experiments for example have linked cortical remapping to experience-dependent plasticity and long-term potentiation (LTP) (Glazewski et al. 1996; Hardingham et al. 2008; Margolis et al. 2012). The pairing of a sensory stimulus with artificially evoked neuronal spikes has been shown to induce a plasticity associated with receptive field dynamics (Jacob et al. 2007; Gambino and Holtmaat 2012; Pawlak et al. 2013; El-Boustani et al. 2018). High frequent and tetanic sensory stimulation can increase sensory-evoked network potentials, which in humans has been demonstrated to lower the response threshold to sensory stimuli (Mégevand et al. 2009; Frenkel et al. 2006; Clapp et al. 2005; Kalisch, Tegenthoff, and Dinse 2008; Marzoll et al. 2022; Lengali et al. 2021; Han et al. 2015; Sanders et al. 2018). Whereas passive daily sensory experience can cause a reduction (‘habituation”) in representation of the experienced sensory stimuli by cortical pyramidal neurons (Kato, Gillet, and Isaacson 2015).

When exploring their environment rodents actively move their whiskers over surfaces and objects in rhythmic sweeps ranging in frequencies from 5-15 Hz, which has been equated to digital palpation and microsaccades in primates (Carvell and Simons 1990; Wolfe et al. 2008). Neuronal membrane potentials and spiking in the S1 are modulated in synchrony with whisking frequency (Fee, Mitra, and Kleinfeld 1997; Crochet and Petersen 2006). Therefore, passive sensory stimulation within natural frequencies may reveal key physiological mechanisms that underpin experience-dependent plasticity. It was previously shown that, under anesthesia, a brief (1min) period of rhythmic whisker stimulation (RWS) at 8Hz enhances whisker-evoked local field potentials and evokes LTP in layer L2/3 pyramidal neurons in S1 (Gambino et al. 2014; Mégevand et al. 2009). This sensory-evoked LTP can be elicited in the absence of somatic spikes and is driven by N-methyl-D-aspartate receptor (NMDAR) -mediated long-lasting depolarizations that remain subthreshold (Gambino et al. 2014). It remains unclear in awake conditions whether RWS leads to changes in cortical population-wide activity, or how this impacts responsivity of individual neurons to subsequent sensory stimulation, over what timescales, and if this is driven directly by whisker input (Gambino et al. 2014; Williams and Holtmaat 2019; Mégevand et al. 2009). Furthermore, it has also been shown that this plasticity may require the activation of a disinhibitory gating circuit motif that involves vasoactive-intestinal-peptide-expressing (VIP) interneuron activity (Williams and Holtmaat 2019). It is not known if the activity of VIP interneurons is modulated during RWS.

To assess to what extent neuronal population activity and stimulus representation in the cortex is impacted by RWS we monitored whisker-evoked calcium (Ca^2+^) signals in L2/3 neurons and VIP interneurons in S1 for hours upon RWS in awake mice. In S1 whiskers are functionally represented in the barrel cortex, and neurons in each barrel-related cortical column (barrel column hereafter) responding best to a single whisker. We found that RWS produced a potentiation of whisker-evoked responses in many L2/3 neurons (96%) that reside in the parent barrel column of the stimulated whisker – termed the principal whisker (PW), and which display low or moderate responses under baseline conditions. When a distant whisker – termed control whisker (CW) was used for RWS, a very small (4%) high responding neuronal population significantly decreased their subsequent PW-evoked activity. We also found that RWS of the PW altered its representation in the parent column by preferentially recruiting or retaining active neurons to the responsive pool, whereas RWS of the CW tended to suppress responsivity. The potentiation of sensory-evoked activity lasted for at least 1h on average. VIP interneurons displayed a sustained nonselective increase in activity during the RWS period, but their PW-evoked responses after RWS were not potentiated. These findings suggest that repetitive whisker stimulation, within the range of frequencies at which mice sense objects, causes a selective potentiation of sensory-evoked responses and recruits’ neurons to respond to subsequent sensory stimulation. Moreover, this may be supported by a whisker-non-selective VIP interneuron-mediated disinhibitory mechanism.

## METHODS

### Experimental Model and Subject Detail

*Animals.* 5-7-week-old C57BL/6J male mice (Janvier Labs) or Vip-IRES-cre (*Vip^tm1(cre)Zjh^*/J; The Jackson Laboratory, RRID: IMSR_JAX:010908) (Taniguchi et al. 2011) were grouped housed on a 12h light cycle with littermates. All procedures were conducted in accordance with the guidelines of the Federal Food Safety and Veterinary Office of Switzerland and in agreement with the veterinary office of the Canton of Geneva (license numbers GE/28/14, GE/61/17, GE/74/18, and GE253).

### Method Details

*Surgery and virus injections.* Stereotaxic injections of adeno-associated viral (AAV) vectors were carried out on 6-week-old male C57BL/6 mice. A mix of O_2_ and 4% isoflurane at ∼0.4l min ^-1^ was used to induce anesthesia followed by an intraperitoneal injection of MMF solution, consisting of 0.2mg kg^-1^ medetomidine (Domitor, Orion Pharma), 5mg kg^-1^ midazolam (Dormicum, Roche), 0.05mg kg^-1^ fentanyl (Fentanyl, Sintetica) diluted in sterile 0.9% NaCl. AAV2-CAG-GCaMP6s-WPRE-SV40 (U Penn Vector Core, RRID: Addgene_100844, 100 nl) or AAV1-hSyn-mRuby2-GSG-P2A-GCaMP6s-WPRE-pA (addgene, RRID: Addgene_50942; 100 nl) was delivered to L2/3 of the right barrel cortex at the approximate location of the C2 barrel column (1.4mm posterior, 3.5mm lateral from bregma, 300mm below the pia) (Rose et al. 2016). For targeting VIP interneurons AAV1-CAG-flex-mRuby-P2AGCaMP6s-WPRE-pA (addgene, RRID:Addgene_68717) was injected into the VIP-IRES-cre mouse line and was repeated 3 times around the same area (3x50nL) (Rose et al. 2016). For long-term *in vivo* Ca^2+^ imaging a 3-mm diameter cranial window was implanted, as described previously (Holtmaat et al. 2009).

Two weeks after surgery, the barrel columns were mapped using intrinsic optical imaging and the intrinsic optical signal (iOS) was used to confirm the barrel specific location of GCaMP6s expression (Fig. 1A). Anaesthesia was induced using isoflurane (4% with ∼0.4 L.min^-1^ O_2_) and then continued using an intraperitoneal injection of MM consisting of 0.2mg kg^-1^ Medetomidine (Domitor, Orion Pharma), 5mg kg^-1^ Midazolam (Dormicum, Roche) diluted in sterile 0.9% NaCl. Body temperature was maintained at 37°C using a feedback-controlled heating pad.

**Figure 1:**
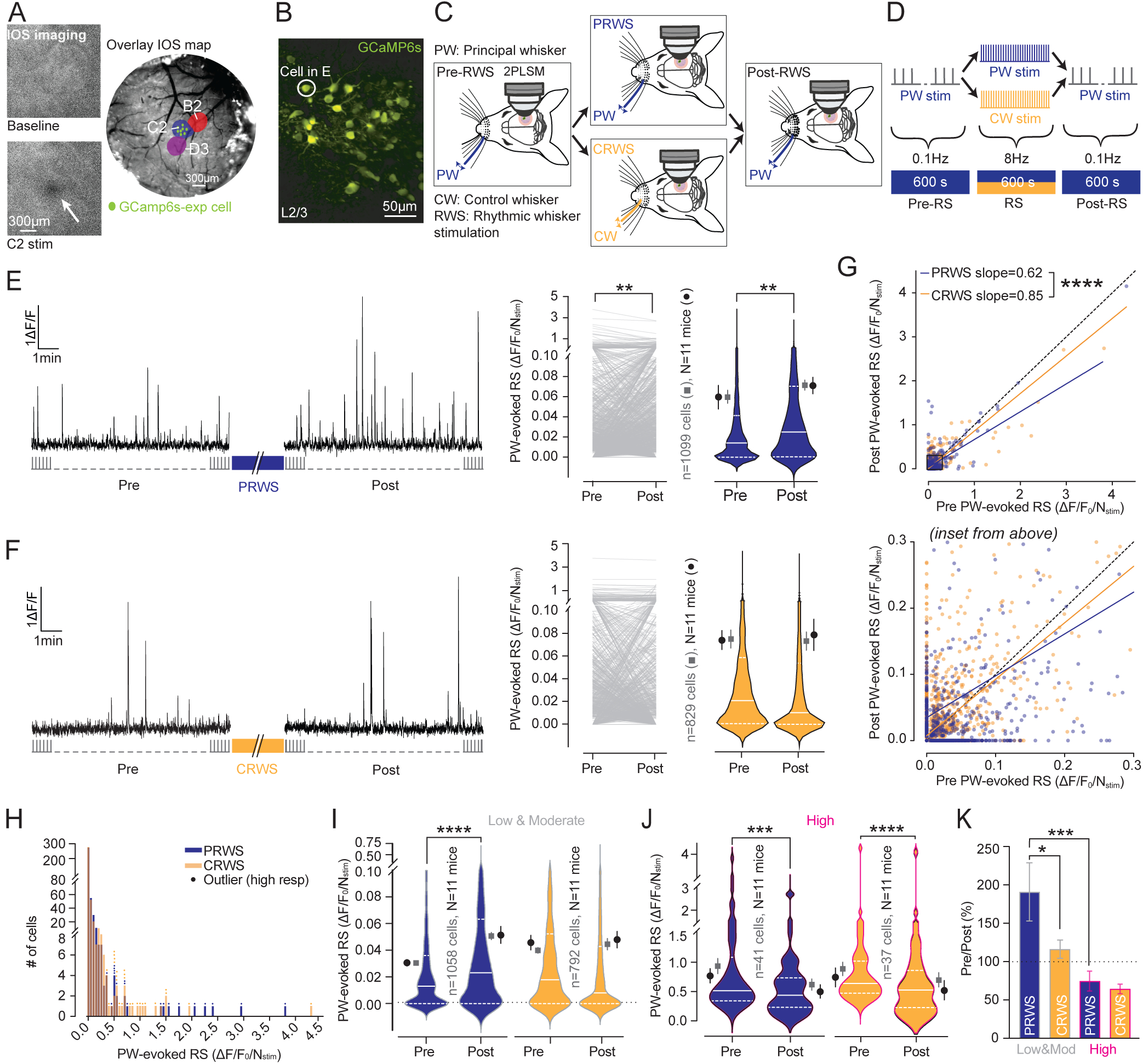
PRWS potentiates whisker-evoked responses in L2/3 neurons. **(A)** Left, examples of averaged baseline & stimulus-related raw iOS images, evoked by one train of whisker deflections. Right, example barrel map overlayed over a brightfield image of the blood vessels. Green dots represent the location of GCaMP6s-expressing cells in the C2 barrel column. **(B)** Average 2PLSM image of GCaMP6s-expressing neurons. **(C)** Experimental design: the PW, which corresponds to the barrel column containing the GCaMP6s-expressing cells, is always used to read out the sensory stimulus-evoked response. The PW for PRWS (blue) or a control far-away whisker (CW) for CRWS (orange) is stimulated during rhythmic whisker stimulation (RWS, 8Hz, 10min). **(D)** Experimental protocol: the PW is stimulated at 0.1 Hz for 10 min pre- and post-RWS. RWS (8 Hz, 10 min) is performed on either the PW (PRWS, blue) or a far-away CW (CRWS, orange). Whisker movement index during the stimulus protocol can be found in Figure 1-1. **(E & F)** Left, example trace of the GCaMP6s fluorescence, in response to PW stimulation (0.1 Hz, 10 min) pre- and post-PRWS (**E**) or CRWS **(F)**. The signals in **E** are from the cell circled in **B**. Right, the PW-evoked response strength (RS, amplitude X whisker-evoked signal probability, (ΔF/F_0_)/Nstim) pre- and post-PRWS (**E**, n=1099 cells, **P=0.002, N=11 mice, P=0.5, full descriptive statistics can be found in Table 1-1) or CRWS (**F**, n=829 cells, P=0.4; N=11 mice, P=0.6). Grey lines, paired responses. Violin plots depict median (solid) and quartiles (dotted) bars. Squares, the mean over cells (±SEM). Circles, the mean over mice (±SEM). **(G)** Pre- versus post-RWS RS ((ΔF/F_0_)/Nstim) with the simple linear regression for PRWS (blue, n=1099 cells) and CRWS (orange, n=829 cells). Comparing slopes (PRWS=0.62±0.014, CRWS=0.85±0.019, F=92.9, DFn=1, DFd=1924, ****P<0.0001). **(H)** Frequency distribution of the pre-RWS RS for PRWS & CRWS, bin size 0.01(ΔF/F_0_)/Nstim. High responders (resp) were identified as outliers (dots above, Iterative Grubb’s outlier test, α=0.0001). **(I & J)** Violin plot of the RS pre- and post-PRWS and CRWS, for low & moderate responders **(I**, PRWS, n=1058 cells, ****P<0.0001; N=11 mice, P=0.006; CRWS, n=792 cells, P=0.3; N=11 mice, P=0.7), and for high responders (**J**, PRWS, paired t-test, n=41 cells, ***P=0.0008; N=11 mice, P=0.04; CRWS, n=37 cells, ****P<0.0001; N=11 mice, P=0.0001). **(K)** The pre-RWS/post-RWS ratio (in %) for PRWS and CRWS low & moderate, and high responders (mixed effects model, N=11 mice, P=0.0004; multiple comparisons: PRWS low & moderate vs high, ***P=0.0004; CRWS low & moderate vs high, P=0.1; Low & moderate PRWS vs CRWS, *P=0.018; High PRWS vs CRWS, P=0.7).

To illuminate the cortical surface through the cranial window, a light guide system with a 700 nm (bandwidth of 20 nm) interference filter and a stable 100-W halogen light source were utilized. Images were acquired using the Imager 3001F (Optical Imaging, Mountainside, NJ) equipped with a large spatial 256*256 array, a fast readout, and a low read noise charge coupled device (CCD) camera. The size of the imaged area was adjusted by using a combination of two lenses with different focal distances (Nikon 50mm, bottom lens, 135mm, upper lens, f=2.0; total magnification 2.7). The CCD camera was focused on a plane 300µm below the skull surface. Mapping was started by first inserting the C2 whisker in a glass capillary attached to a piezo actuator (PL-140.11 bender controlled by an E-650 driver; Physik Instrumente). Whisker stimulations were triggered by a pulse stimulator (Master-8, A.M.P.I.). In a typical session, 10 trials were collected per stimulus. Intrinsic signals were acquired at 10Hz for 5s (50 frames, 100ms per frame). Each trial lasted 5s and consisted of a 1s pre-stimulus baseline period (frames 1-10), followed by a 1s stimulus period (11-20), during which the whisker was deflected (1s at 8Hz), and then by a 3s post-stimulus period (frames 21-50). Inter-trial intervals lasted 15 to 20s. Responses were visualized by dividing the stimulus signal by the baseline signal, using the built-in Imager 3001F analysis program (Optical Imaging, Mountainside, NJ). If needed, other whiskers were subsequently stimulated to generate a map. An image of the surface vascular pattern was taken using green light (546 nm interference filter) and superimposed onto the intrinsic signal image which created reference image to the barrel map. This reference image was used later to select the appropriate PW for the visualized fluorescent neurons as well as a CW that was minimally 2 rows and 2 arcs away. After this procedure, a metal post was glued onto the head cap laterally to the window using dental cement.

*Mice habituation.* After intrinsic imaging, mice were handled for 20-30 minutes each day. On the second day they were placed in the imaging holder for 30s, with this habituation time increasing over the subsequent days.

*Rhythmic Whisker Stimulation (RWS) protocol.* The whisker which produced an iOS that most strongly overlapped with the location of the fluorescently labeled neurons was identified as the PW (e.g. C2 in Fig. 1A,B). Mice were briefly anesthetized using isofluorane (4% with ∼0.5l min -1 O_2_) to identify and mark the PW with colored nail polish. RWS was either performed using the PW (PRWS), or using a CW (CRWS) that was minimally 2 rows and 2 arcs away from the PW (Fig. 1B,C). The CW, therefore, had a poor anatomical and functional connectivity with the PW home barrel column. The control protocol (Pre, CRWS, and Post) or test protocol (Pre, PRWS, and Post) was applied with approximately 10 days in between. For all experiments pre and post RWS the whisker was deflected back and forth (5 deflections lasting each 45ms @ 20Hz) for 10min at a frequency of 0.1Hz. The whiskers were deflected with a piezoelectric ceramic elements attached to a glass pipette 4 mm away from the skin. The voltage applied to the ceramic was set to evoke a whisker displacement of 0.6 mm with a ramp of 7– 8ms. Different whiskers were independently deflected by different piezoelectric elements, one for the PW and one for CW simulation. For PRWS the PW and for CRWS the CW was deflected back and forth for 10min at a frequency of 8Hz.

*In vivo 2-photon laser-scanning microscopy (2PLSM) imaging*. Across all experiments, 2PLSM imaging of Ca^2+^ signals in GCaMP6s-labeled L2/3 neurons was performed in the home barrel column of the earlier identified PW, and the same field of view was imaged for both experimental conditions for a single mouse. To return to the same field of view over multiple imaging sessions, and to ensure accurate whisker deflection, a brightfield image of the blood vessel pattern directly above the center of the fluorescence expression was taken as a reference guide.

Ca^2+^ imaging was performed using custom built 2PLSMs (https://www.janelia.org/node/46028), controlled by Scanimage 2016b44 (http://www.scanimage.org). Excitation light was provided by Ti:Sapphire lasers (Coherent) tuned to λ=910nm for GCaMP6s signal alone and λ=980 nm for imaging of mRuby2 and GCamp6s. For detection, we used GaAsP photomultiplier tubes (10770PB-40, Hamamatsu), a *16x 0.8NA* microscope objective (Olympus or Nikon, CFl75). When required, mRuby2 and GCaMP6s signals were separated with a dichroic mirror (565dcxr, Chroma) and emission filters (ET620/60 m and ET525/50 m, respectively, Chroma). For imaging L2/3 neurons with constitutive GCaMP6s, the size of the field of view ranged from 187 μm x 187 μm to 375 μm × 375 μm, while pixel size ranged from 0.7 to 1.4, and the imaging speed was set at 3.91Hz (256 lines, 1ms per line). For extended depth of field imaging, the 2PLSM was equipped with an 8-kHz resonant scanner and Axicon setup for Bessel beam generation. Fast volumetric imaging was performed at 10 or 11.5 Hz using a piezo z-scanner (P-725 PIFOC, Physik Instrumente) to move the objective over the z-axis (Meng, Zhang, and Ji 2023). Each acquisition volume consisted of 2 planes P1, that was 80-120µm from pia, and P2 that was 100μm below (180-220µm respectively) of 400 x 400 μm (512 x 256 pixels). This allowed for post-hoc z-motion correction of brain motion artefacts induced by movement. Mice were monitored using an infrared camera across all imaging sessions.

*Data analysis.* Images were processed using custom-written MATLAB scripts and ImageJ/Fiji (http://rsbweb.nih.gov/ij/). Motion correction was performed using a custom strategy based on the cross correlation of the first image compared to subsequent images. Remaining movements were addressed by calculating in Fiji’s correlation plugin another cross correlation 2D graph. From this, the mean value of the cross correlation (between the first image and image 1,2,3…n) was calculated. If a value was outside the mean ±2 SD, it was labelled as movement, and Ca^2+^ signals during this period were not taken into account. For depth of field imaging, lateral and axial motion corrections were performed and the mRuby2 signal was used as a reference – as previously described using NoRMCorre (Flatiron Institute, Simons Foundation, New York, NY 10010, USA, https://github.com/flatironinstitute/NoRMCorre) (Chéreau et al. 2020; Pnevmatikakis and Giovannucci 2017). Regions of interest (ROIs) were drawn by hand using the GCaMP6s channel, or the mRuby channel when present. Pixels were averaged within each ROI for each image frame. For each ROI, normalized Ca^2+^ traces DF/F_0_ were calculated as (F−F_0_)/F_0_, where F_0_ is the 30^th^ percentile of the individual mean baseline fluorescence signal for the entire recording session (pre, RWS, and post). For each ROI, Ca^2+^ signals were detected in Caltracer3beta, using a fluorescence intensity threshold (0.5 DF/F_0_), amplitude threshold (1DF/F_0_) and rise time threshold (0.256s, http://www.columbia.edu/cu/biology/faculty/yuste/methods.html) (Ayzenshtat et al. 2016). Whisker stimulations across the recording session were aligned with the Caltracer3beta detected Ca^2+^ events and were considered a PW-evoked event if their rise time occurred within 512ms. All other events were considered spontaneous and not included in the analysis.

For each neuron (either pre or post-RWS), we calculated the PW-evoked Ca^2+^ signal probability (*P_S_*), the average PW-evoked Ca^2+^ signal amplitude (*Ā_S_*), and the PW-evoked response strength (*RS* [DF/F_0_/N_stim_]), as follows:

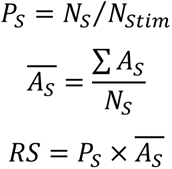

where *N_S_* and *N_Stim_* are the number of PW-evoked Ca^2+^ signals and the total number of stimuli respectively, and *A_S_* is the amplitude of a single evoked Ca^2+^ signal. For the mean fluorescence intensity during RWS (DF/F_0_) was calculated and the Ca^2+^ fluorescence intensity was integrated over the baseline (20s before start of RWS) and during the RWS period (20s after the start of RWS). All data are reported as mean± standard error of the mean (SEM) per cell and per mouse (**See Table 1-1. Descriptive Statistics**). Stats we performed per mouse and cell to test the robustness of the effect.

*Whisker movement index.* Whisker movements were tracked on 4 mice both pre- and post-RWS (separate set from the Ca^2+^ imaging dataset). The habituation and RWS protocol were as above. Whisker pads on either side of the snout were imaged from below at 112Hz using a Point Grey Research USB 2.0 CCD Digital Camera (Model 00-00100-08200) and Streampix 9 software. To extract whisker movements, we used a custom-written MATLAB script. Here, ROIs were drawn on whiskers ipsi- and contra-lateral to the capillary. For these ROIs, we first calculated the correlation between each frame and the average image of the entire movie (Fig. 1-1A). Next, we computed the absolute derivative of this cross-correlation. We calculate the absolute derivative of the cross correlation to obtain a value for the whisker movement, i.e., the movement index (MI) with arbitrary units (a.u.). Lastly, a Savitzky-Golay filter was applied to the trace to smoothen it, and thus each point/value corresponds to 1 frame in the movie (Fig. 1-1B). To compare the difference in whisking before and after each stimulation, we calculated the average MI over 2s (224 frames) before the start and after the end of the stimulation (Fig. 1-1C).

*Immunohistochemistry.* At the end of the experiment, animals were anesthetized and fixed using transcardial perfusion of 4% paraformaldehyde (PFA) in saline. The brains were left in 4% PFA overnight (4 °C) for further fixation. Coronal brain sections (50μm thickness) were obtained using a vibratome (Leica VT1200S; Leica Microsystems, Vienna, Austria) and initially stored in PBS. Fluorescence microscopy was used to confirm the injection site. For immunohistochemical detection and quantification of VIP interneurons, slices were then incubated for 1h, free floating in a blocking solution of PBS (pH 7.4) containing 0.3% Triton and 1% Bovine Serum Albumin (BSA). After blocking, slices were incubated overnight in blocking solution containing primary antibody (VIP, rabbit polyclonal IgG, Immunostar, Cat#:20077, RRID: AB_572270) at a 1:500 dilution(Williams and Holtmaat 2019). Slices were washed 4 times for 10mins each in PBS and 5% BSA at room temperature. They were then incubated for 1h in PBS solution containing 1% BSA and the appropriate fluorescence conjugated secondary antibodies (1:400, Goat anti-Rat IgG (H+L) Cross-Adsorbed Secondary Antibody, Alexa Fluor 647, Thermo Fisher Scientific, Cat#: A32733, RRID: AB_2633282). Finally, slices were washed 4 times in PBS at room temperature. Cell nuclei were stained using Hoechst 33342 (Invitrogen, Cat#: H1399, RRID: AB_10626776) diluted 1:5000 in PBS and added for 20 mins. Lastly, slices were washed 4x in PBS and placed onto glass slides.

Images were generated using a confocal laser-scanning fluorescence microscope (Nikon A1 R) at 20x magnification. Fluorescence intensity was measured by delineating the edges of all visible cells using ImageJ software and by calculating mean fluorescence in these ROIs. To avoid false-positives, two controls were performed. First, images were taken in an area adjacent to injection area (i.e. cells that were not visibly expressing mRuby). ROIs were drawn around anti-VIP positive cells, and fluorescence intensity in the red channel was quantified. Second, images were taken within the injection area in sections on which only the secondary antibody Alexa 647 was applied. ROIs were drawn around cells, and fluorescence intensity in the green channel was quantified. Each of these quantifications yielded a mean fluorescence – 2SD, which was subsequently used as the lower-limit on which we based the overlap estimate (i.e., no. Of true positives/total no.). Intensities of the experimental cells below these limits were considered as false positive in either channel.

### Experimental Design and Statistical Analysis

For all experiments, n equals the number of cells and N equals number of mice. The statistics were performed over cells (grey box in figures, stats reported in the text and figure legends) and over mice (black circles in figures, and stats reported in figure legends and extended data Table 1-1). All statistics were performed, and graphs were created using Prism 9 or 10 (GraphPad Software, LLC). For all figures, significance levels were denoted as *P < 0.05, **P < 0.01, ***P < 0.001, and ****P < 0.0001 and asterisks were reported per cell in the figures. No statistical methods were used to estimate sample sizes. A paired t-test was performed unless otherwise noted. All comparison tests were performed two-sided. All data are reported as mean± standard error of the mean (SEM).

## RESULTS

### RWS modulates whisker-evoked activity of L2/3 cortical neurons

To monitor whisker-evoked activity of L2/3 neurons in the barrel cortex of S1, we expressed the genetically encoded Ca^2+^ sensor GCaMP6s using adeno-associated viral vectors (AAV2-CAG-GCaMP6s or AAV1hsyn-mRuby2-GSG-P2A-GCaMP6s). Single cell Ca^2+^ signals were recorded using two-photon laser scanning microscopy (2PLSM) through a chronically implanted cranial window (Fig. 1B). The location of GCaMP6s-expressing neurons relative to the whisker representations in the barrel cortex was determined using intrinsic optical signal imaging (iOS see methods; Fig. 1A). The whisker which produced an iOS that most strongly overlapped with the location of the fluorescently labelled neurons was identified as their principal whisker (PW, e.g. C2 in Fig. 1A).

We monitored the stimulus-evoked Ca^2+^ signals, always upon stimulation of the PW (10 min, 0.1 Hz) pre- and post-rhythmic whisker stimulation (RWS, Fig. 1C, D). For RWS (10 mins, 8Hz) we used either the PW (PRWS) or a control whisker (CW; CRWS), which was minimally 2 rows and 2 arcs away from the PW as defined by the iOS map (Fig. 1A, C, D). The CW, therefore, had poor anatomical and functional connectivity with the PW home barrel column.

Extracted Ca^2+^ signals were classified as PW-evoked events when their onset occurred within 512 milliseconds (ms) after the start of a PW stimulus. The stimulation response window was determined by acquisition frame rates and typical GCaMP6s response kinetics (Chen et al. 2013). For each neuron, we calculated the PW-evoked response strength (RS; hereafter simply ‘response strength’) as a product (RS [DF/F_0_/N_stim_] = *Ā_S_* x *P_S_*) of the Ca^2+^ signal’s amplitude (*Ā_S_)* and whisker evoked signal probability (*P_S_)*.

We found that PRWS increased the mean response strength of the whole L2/3 neuronal population (Fig. 1E, pre-PRWS mean±SEM=0.06±0.007, post-PRWS=0.07±0.005, n=1099 cells, P=0.0015) but not when we compared the population averages over mice (N=11 mice, P=0.5; for descriptive statistics over cells (n, grey square) and mice (N, black circle, see extended data Table 1-1). Upon CRWS, however, the mean response strength remained unchanged (Fig. 1F, pre-CRWS=0.076±0.008, post-CRWS=0.072±0.008, n=829 cells, P=0.4; N=11 mice, P=0.6). The pre-PRWS response strength was somewhat lower, but not significantly different from the pre-CRWS (P=0.1), which had likely resulted from different fields of view for the two paradigms.

When we plotted the pre versus post-RWS response strength and ran a simple linear regression analysis on the population data, we found for CRWS the slope only slightly deviated from the identity line (slope=0.85; Fig. 1G), whereas for the PRWS the slope was significantly lower (slope=0.62, P<0.0001). For PRWS the linear regression line crossed the identity line, indicating that the neurons with a low baseline response strength were more likely to be potentiated, whereas those that initially showed a high response strength tended to be depressed. For CRWS, the neurons with a high baseline response strength also tended to be depressed, but those with a low response strength were not potentiated.

Since the linear regression analysis suggested that the plasticity was dependent upon the baseline response strength, we looked more in detail at the baseline properties. The vast majority had a relatively low or moderate (96%, n=1058 cells) response strength, whereas a small group of neurons showed a relatively high response strength. When we tested the mean pre-RWS response strength for significant outliers (Fig. 1H, Iterative Grubb’s outlier test, α=0.0001), we found that 4% of the population exhibited an excessively high mean pre-RWS response strength (pre-PRWS=0.87±0.13, n=41 cells, pre-CRWS=0.72±0.06, n=37 cells, N=11 mice). We categorized those as high responders (Fig. 1H) (Margolis et al. 2012; Crochet et al. 2011). All mice had high responders in their L2/3 neuronal population.

When we analyzed the response strength for groups separately, we found that for the low & moderate responders, the PRWS increased the mean response strength (Fig. 1I, pre-PRWS mean±SEM=0.03±0.001 DF/F_0_/N_stim_, post-PRWS=0.05±0.003, n=1058 cells, P=0.0015), but not the CRWS (Fig. 1I, pre-CRWS=0.04±0.002, post-CRWS=0.045±0.004, n=792 cells, P=0.3). In contrast, for the high responders, we found a significant decrease in response strength for both PRWS (Fig. 1J, pre-PRWS=0.9±0.13, post-PRWS=0.6±0.09, n=41 cells, P=0.0008) and CRWS (pre-CRWS=0.84±0.11, post-CRWS=0.66±0.12, n=37 cells, P<0.0001).

Further analysis revealed a significant difference in the size of the change in response strength between low & moderate and high responders overall (Fig. 1K, mixed-effects model, P=0.0004). For low & moderate responders, the change was significantly higher upon PRWS as compared to CRWS (190.0±37.8% vs. 116.3±11.9%, Uncorrected Fisher’s LSD, P=0.018). Moreover, the change upon PRWS for low & moderate responders was significantly higher as compared to high responders (190.0±37.8% vs. 74.7±12.9%, Uncorrected Fisher’s LSD, P=0.0004). There was no difference between the two groups for CRWS (74.7±12.9% vs. 64.2±6.24%, Uncorrected Fishers LSD, P=0.7). Overall, this suggests that PRWS induces a potentiation of subsequent PW-evoked responses for the majority of the imaged L2/3 neurons, dependent on the baseline whisker-evoked response strength. The low & moderate responders increase their response, whereas the high responders tend to lower their response. This bears similarities to experience-dependent plasticity effects upon trimming all but one whisker, upon which low responders increase and high responders decrease their responsiveness to the spared whisker (Margolis et al. 2012).

To exclude the possibility that the above effects were merely driven by changes in overall whisking rates pre- and post-RWS or due to reactive whisking upon the stimulus, we performed an additional experiment in which we monitored whisker movements on the ipsilateral (i.e. capillary tube side) or the contralateral side during the same protocol. The mice (n=4) were habituated similarly to above, but now whiskers were imaged using a CCD camera (112 Hz frame rates) positioned below the whisker pads (Fig. 1-1A). Using custom software, a whisker movement index (MI) was calculated 2s before and after each whisker stimulus, pre- and post-RWS. We found no difference in the average MI pre versus post-RWS (Fig. 1-1B, ipsi pre=1.0, post=0.98±0.03; contra pre=1.19±0.11, post=1.12±0.09, one-way ANOVA, P=0.24), which strongly suggests that overall whisking rates were not affected by the protocol. Furthermore, for all stimuli the before- and after-stimulus MI were highly variable and only moderately correlated. Furthermore, the average before- and after-stimulus MIs in both the pre- and post-RWS periods were not statistically different on either side of the snout, indicating that the whisker-stimulus did on average not elicit whisking bouts (Fig. 1-1C)

Together, these data suggest that the changes in Ca^2+^ signals which we had observed in our earlier experiments were not due to alterations in whisking behavior, which supports our conclusion that they were primarily related to adaptations in sensory-cortical synaptic pathways.

### RWS heterogeneously potentiates, recruits, and suppresses sensory-evoked responses

Cortical neuronal populations can show a remarkable heterogeneity in sensory-evoked activity (Sato et al. 2007; Brecht, Roth, and Sakmann 2003; Crochet and Petersen 2006; Kerr, Greenberg, and Helmchen 2005; Margolis et al. 2012). Consistent with these findings, we had identified two groups of neurons, those with low & moderate and high responding profiles (see above). In addition, based on the changes in response strength that we observed upon RWS, we could subdivide the low & moderate responders into subgroups. We found that nearly half of the imaged neuronal population showed one or more PW-evoked Ca^2+^ signals pre- and post-RWS and were therefore termed as ‘persistent’ cells (Fig. 2A, D). The remaining neurons lacked PW-evoked Ca^2+^ signals pre- and/ or post-RWS. This group was subdivided into those that lacked signals pre-RWS but displayed them at any time point post-RWS – termed ‘recruited’ cells (Fig. 2B, D); those that displayed PW-evoked signals pre- RWS but not at any time point post-RWS – termed ‘suppressed’ cells (Fig. 2C, D); and finally, cells that showed no events at any time point – termed ‘no-response’ cells (Fig. 2D). We found that the ratios of persistent, recruited, suppressed, no response-cells, and high responders were significantly different between PRWS and CRWS (Chi square test, P<0.0001). For PRWS we found more persistent (PRWS: 42%, CRWS: 34%) or recruited (PRWS: 28%, CRWS: 16%), and less suppressed (PRWS: 19%, CRWS: 31%) or no-response (PRWS: 7%, CRWS 15%) cells than for CRWS. High responders made up 4% of the total population for both PRWS and CRWS.

**Figure 2:**
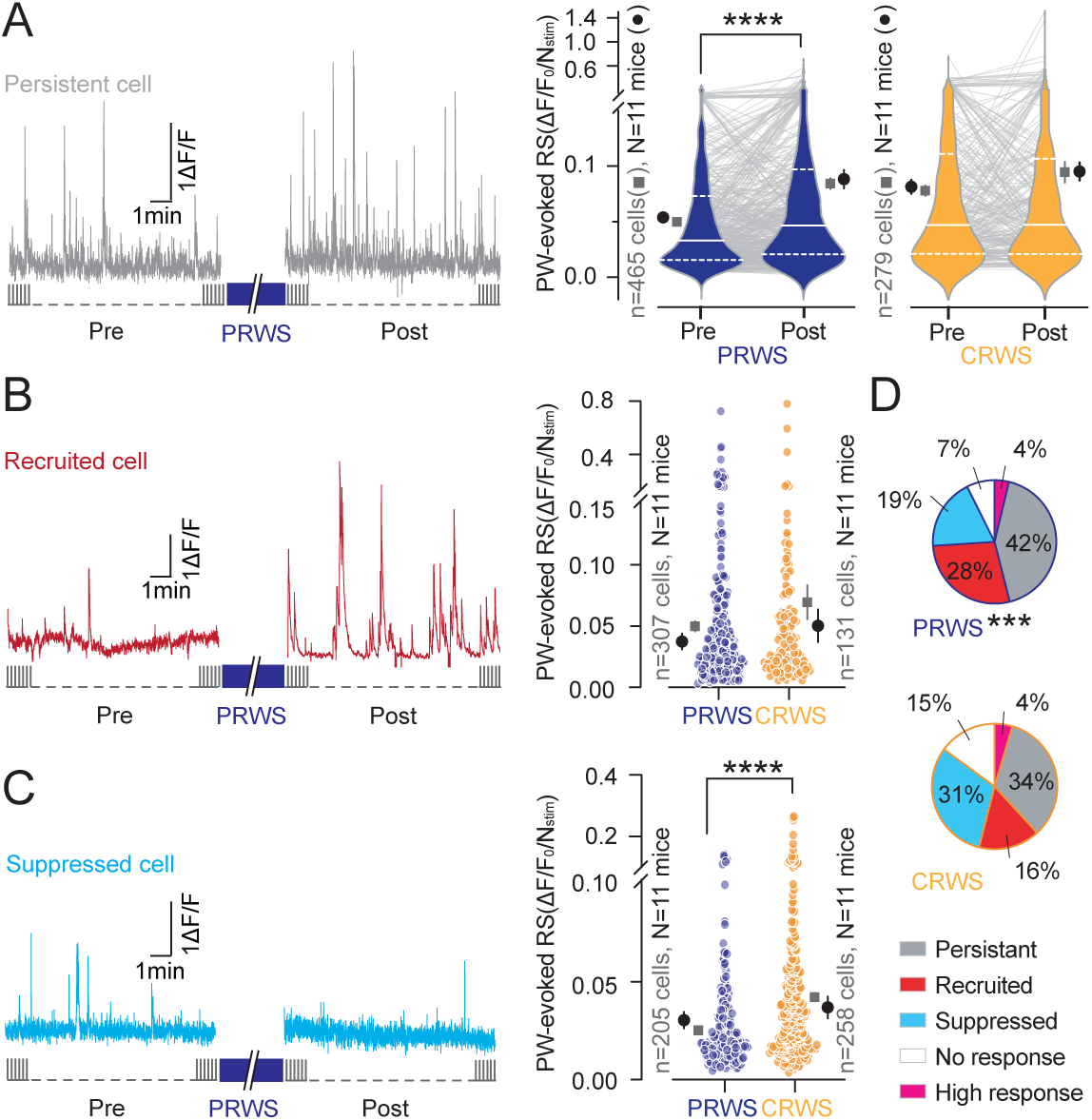
PRWS recruits L2/3 neurons to the active pool. **(A)** Left, example trace of GCaMP6s fluorescence from a neuron showing persistent PW-evoked responses (0.1 Hz, 10 min) pre- and post-PRWS. Right, violin and pairwise representation of pre- and post-PRWS (n=465 cells, ****P<0.0001; N=11 mice, P=0.003) or CRWS (Paired t test, n=279 cells, P=0.06; N=11 mice, P=0.2). **(B&C)** Left, example trace of GCaMP6s fluorescence from neurons of which responses were recruited **(B)** or suppressed **(C)** post-PRWS. **(B)** Right, the mean PW-evoked response strength (RS) of recruited neurons, post-PRWS (n=307 cells) & CRWS (Unpaired t test, n=131 cells, P=0.1; N=11 mice, P=0.5). **(C)** Right, the mean response strength of suppressed neurons pre-PRWS (n=205 cells) & CRWS (Unpaired t test, n=258 cells, ****P<0.0001; N=11 mice P=0.037). **(D)** Pie charts with the percentages (%) of persistent (grey), recruited (red), suppressed (blue), no response (white), and high (pink) responders for PRWS (n=1099 cells) & CRWS (n=829 cells, Chi-square=19.8, DF=4, ***P<0.0001).

Consistent with our overall findings, the persistent subpopulation exhibited a significant increase in response strength upon PRWS (Fig. 2A, pre-PRWS=0.05±0.002, post-PRWS=0.08±0.006, n=465, P<0.0001), but not upon CRWS (pre-CRWS=0.076±0.005, post-CRWS=0.09±0.009, n=279 cells, P=0.06). For the recruited subpopulation, the mean post-RWS response strength was similar for both conditions (Fig. 2C, post-PRWS=0.05±0.004, n=307 cells, post-CRWS=0.07±0.001, n=131 cells, Unpaired t test, P=0.14). For the suppressed subpopulation however, the mean pre-RWS response strength was significantly lower for PRWS when compared to the CRWS condition (Fig. 2D, pre-PRWS=0.03±0.002, n=205 cells, pre-CRWS=0.04±0.003, n=258 cells, Unpaired t test, P<0.0001). This suggests that PRWS may prevent suppression of low & moderate responding cells, whereas CRWS fails to keep those cells in the responsive population.

Overall, we found that PRWS potentiates the PW-evoked activity of neurons that persistently but moderately respond to the PW and recruits new neurons to the active pool (for PRWS 28%), while selectively suppressing the activity of neurons that initially responded either very weakly or very strongly to the PW (Fig. 2D, E).

### Longitudinal imaging upon RWS

Next, we tracked how the potentiated responses advanced over time in individual neurons. To this end we used a bicistronic AAV construct that drives the co-expression of GCaMP6s and mRuby in L2/3 neurons (AAV1-hsyn- mRuby2GSG-P2A-GCaMP6s, Fig. 3A, B). Only cells in which the cell filler mRuby was present at all timepoints were analyzed and cells (ROIs) were chosen in the mRuby channel. L2/3 neurons were imaged pre-PRWS or pre-CRWS for 10 min (-10), and at four timepoints post-PRWS or post-CRWS for 10 min (at 10, 60, 120, and 180 min). We did not observe any difference in imaging depth (data now shown; PRWS: 132±16.9μm, CRWS: 138.5±20.5, Unpaired t-test, P=0.5) or mean baseline response strength between the PRWS and CRWS conditions (data not shown; pre-PRWS=0.09±0.02, pre-CRWS=0.10±0.20; Unpaired t-test, P=0.7). Like before, when we separated the 4% high responders from the analysis, we found that the mean PW-evoked response strength for a majority of the neurons was significantly increased at 10 and 60 min post PRWS, and had returned to baseline levels after 120 min (Fig. 3B, Dunnett’s multiple comparisons, -10min: 0.03±0.002 vs, 10: 0.045±0.005, P<0.0001; or 60: 0.04±0.005, P=0.0001; or 120: 0.032±0.003, P=0.51; or 180: 0.034±0.004; P=0.4). This was significantly different from the CRWS condition (PRWS n=382 cells, CRWS n=304 cells; Two-way RM ANOVA,***P=0.0006) for which we found no significant prolonged increase in mean PW-evoked response strength (Dunnett’s multiple comparisons, -10min: 0.034±0.004 vs, 10: 0.034±0.005, P=0.9; or 60: 0.031±0.005, P=0.3; or 120: 0.033±0.004, P=0.7; or 180: 0.032±0.004; P=0.5). When comparing the individual components of response strength, probability, and amplitude, we observed a significant increase in mean PW-evoked signal probability (P_S_) and amplitude (Ā_S_) at 10 and 60 min (Fig. 3C, D). For CRWS condition there was no such increase in both probability (Fig. 3C, Table 1-1, PRWS n=382 cells, CRWS n=304 cells; Two-way RM-ANOVA, P<0.0001) and amplitude (Fig. 3D, Table 1-1, PRWS n=382 cells, CRWS n=304 cells; Two-way RM-ANOVA, P=0.001), resulting in a significant difference between PRWS and CRWS for these parameters as well. In a separate set of experiments, we tested if the potentiation of activity was present at 24 hours post-RWS but did not find a significant difference (Fig. 3E, Two-way RM-ANOVA, P=0.36) for PRWS (n=162 cells, –10min: 0.051±0.016, 24hrs: 0.056±0.012, Sidak’s multiple comparison, P=0.79) as compared to CRWS (n=134 cells pre -10 min=0.041±0.006, 24hrs: 0.056±0.01, Sidak’s multiple comparison, P=0.14). Altogether, this indicates that PRWS drives a whisker-selective potentiation of whisker-evoked responses for ∼1 hour.

**Figure 3:**
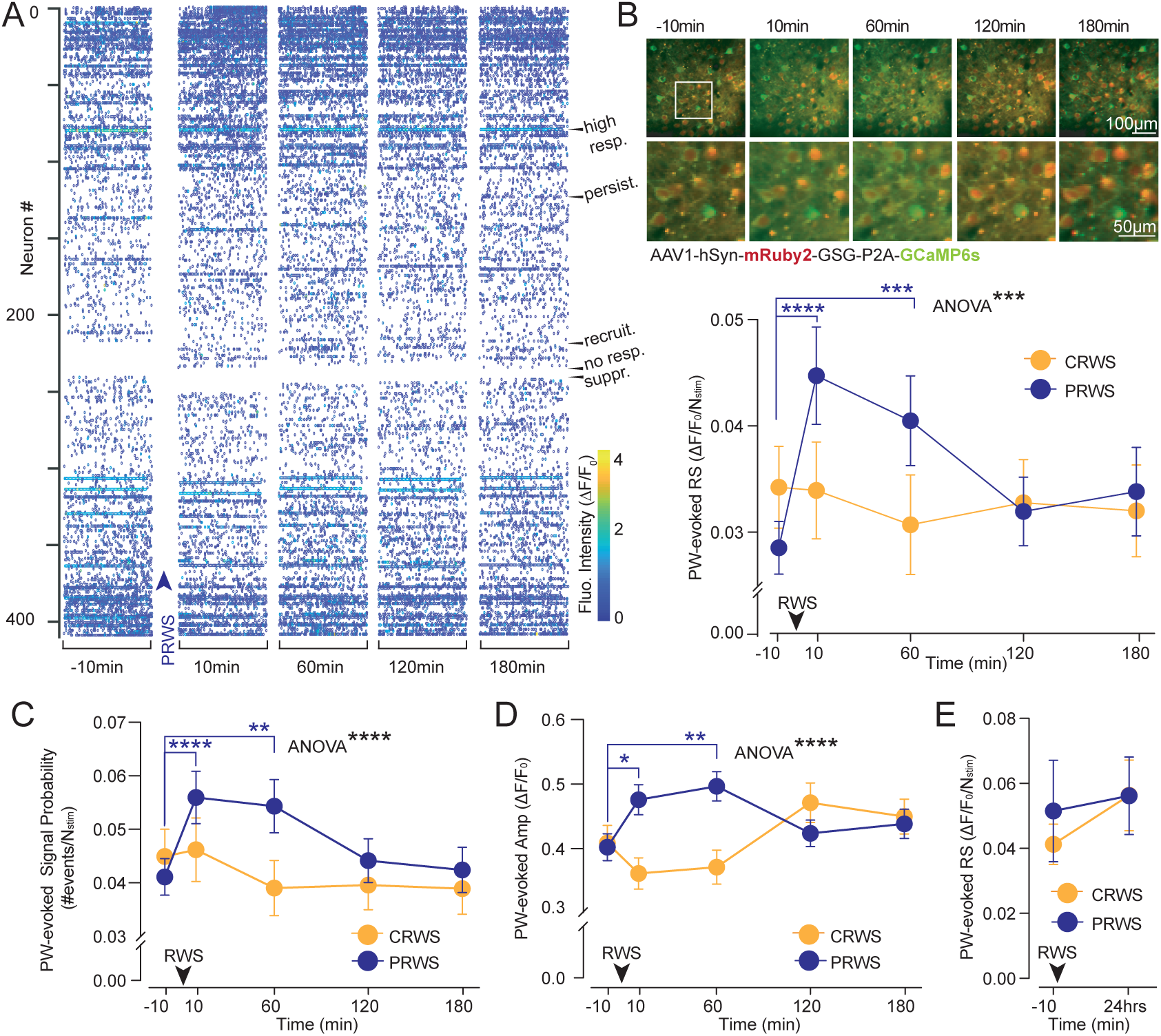
Longitudinal imaging upon RWS. **(A)** Raster plot of GCaMP6s fluorescence intensity (ΔF/F_0_) for PRWS at each acquisition for pre-PRWS (-10 min) and post-PRWS (10, 60, 120, & 180 min). Neurons sorted from top to bottom by decreasing response strength pre- vs post-PRWS (n=410 cells). Arrowheads, examples of the five subpopulations: persistent (persist.), recruited (recruit.), suppressed (suppr.), no response (no resp.), and high (hi resp.) responders. **(B)** Top, example 2PLSM images of neurons expressing AAV1-hSyn-mRubyGSG-P2A-GCaMP6s across the longitudinal experimental protocol -10 min pre-PRWS, & 10, 60, 120, & 180 min post-PRWS. mRuby (red) serves as an activity-independent marker, whereas GCaMP6s (green) reports Ca^2+^ signals upon PW-stimulation. The lower images represent high magnifications of the cells in the square inset on top. **(B)** Bottom, PW-evoked RS pre-PRWS (-10 min) or CRWS and post-PRWS or CRWS (10, 60, 120, & 180 min) (PRWS n=382 cells, or CRWS n=304 cells; Two-way RM ANOVA, ***P=0.0006; N=6 mice, P=0.026). Multiple comparisons for PRWS (Dunnett’s, -10 min vs, 10 ****P<0.0001; or 60 ***P=0.0001). **(C)** PW-evoked Ca^2+^ signal probability (P_S_ [#events/Nstim]) (PRWS n=382 cells, CRWS n=304 cells; Two-way RM ANOVA, ****P<0.0001; N=6 mice, P=0.026). Multiple comparisons for PRWS (Dunnett’s, -10 min vs, 10****P<0.0001; or 60 **P=0.001; or 120 P=0.7; or 180 P=0.99) & CRWS (-10 min vs, 10 P=0.99; or 60, P=0.3; or 120 P=0.2; or 180 P=0.3). **(D)** PW-evoked Ca^2+^ signal amplitudes (Ā_S_ [ΔF/F_0_]) (PRWS n=382 cells, CRWS n=304 cells; Two-way RM ANOVA, ****P<0.0001; N=6, P=0.25). Multiple comparisons for PRWS (Dunnett’s, -10 min vs, 10 *P=0.01; or 60 **P=0.002; or 120 P=0.8; or 180 P=0.4) & CRWS (-10min vs, 10 P=0.2; or 60 P=0.6; or 120 P=0.1; or 180 P=0.5). **(E)** PW-evoked response strength, pre- and 24 hrs post-PRWS or CRWS (PRWS n=162 cells, CRWS n=134 cells; Two-way RM ANOVA, P=0.36; PRWS 3 mice, CRWS 2 mice, P=0.054).

### L2/3 neuronal responses during RWS

Next, we investigated the relationship between the prolonged potentiation of the neurons’ PW-evoked responses and their activity elicited during the RWS period (Fig. 4). We found in a separate set of experiments that PRWS rapidly and significantly increased Ca^2+^ signals when compared to a 20-sec baseline period just before RWS, whereas CRWS did not (Fig. 4A, B; baseline=0.2±0.02, PRWS=0.3±0.02, n=260 cells, Paired t test, p<0.0001; baseline=0.47±0.02, CRWS=0.5±0.03, n=115 cells, Paired t test, P=0.6). We did not, however, find a significant correlation between the increase in Ca^2+^ signals during PRWS and the size of the potentiation, despite observing a significant increase in response strength post PRWS within this subset of experiments (Fig. 4C, Pearson r=-0.06, P=0.5; inset, pre=0.03±0.002 & post=0.03±0.002, Paired t test, P=0.03). We did on the other hand, find a significant inverse correlation between the mean baseline response strength and the increase in Ca^2+^ signal during PRWS (Fig. 4D; Pearson r correlation, n=115 cells, r=0.35, P=0.0001, simple linear regression, slope=-0.008). Therefore, low & moderate responders displayed the largest increase in activity during the PRWS period. Altogether, this suggests that that the potentiation of low & moderate responders may indeed depend on the levels of activity elicited by PRWS.

**Figure 4:**
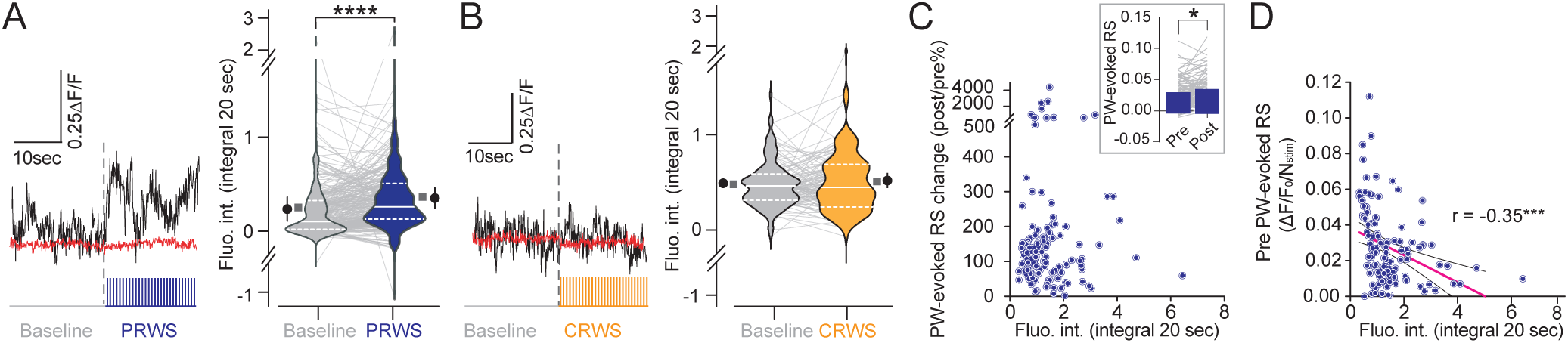
RWS selectively activates L2/3 neurons. **(A, B, Left)** Example traces of GCaMP6s (black) or mRuby fluorescence (red) for a 20-sec baseline before and during PRWS or CRWS. Violin and pairwise representation of fluorescence intensities (integrated over 20s) (violin plot median: white bar, quartiles: dotted bars) baseline vs PRWS (n=290 cells, Paired t test, ****P<0.0001; N=3 mice, P=0.027) or CRWS (n=115 cells, Paired t test, P=0.6; N=3 mice, P=0.8). **(C)** Fluorescence intensity (integrated over 20s) during PRWS vs PW-evoked response strength change (post/pre), (n=115 cells, Pearson r correlation, r=0.0002, P=1.0). Inset, pre- and post-PRWS response strength (n=115 cells, pre=0.027±0.002, post=0.031±0.002, Paired t test, P=0.027). **(D)** Fluorescence intensity during PRWS (integrated over 20s) vs pre-PRWS. Pink line simple linear regression, black dotted lines 95% confidence intervals. (n=115 cells, Pearson r correlation, r=-0.35, ***P=0.0001; simple linear regression, slope=-0.008, non-zero? P=0.0001).

### VIP interneuron activity increases during RWS

We have previously shown that RWS evoked synaptic LTP in L2/3 neurons is gated by a disinhibitory circuit motif that is dependent upon VIP interneuron activity (Williams and Holtmaat 2019). This prompted the hypothesis that the potentiation of L2/3 neuronal sensory-evoked responses may be facilitated by increased VIP interneuron activity during RWS. To assess the activity of VIP interneurons before, during, and after RWS, we injected AAV1-CAG-flex-mRuby-P2A-GCaMP6s into a VIP IRES-cre transgenic mouse line (Fig. 5A). Post-hoc immunostaining analysis confirmed that all mRuby-expressing cells were anti-VIP positive (Fig. 5A, n=199 cells, 81.41% of the cells were Anti-VIP and GCaMP6s-positive, 18.59% were only Anti-VIP-positive, and 0% were only GCaMP6s-positive, data not shown) (Taniguchi et al. 2011). As interneurons make up only ∼20% of the cortical neuronal population, of which ∼13% are VIP interneurons, we employed extended depth of field imaging to capture the activity of a high number of cells per imaging session (Markram et al. 2004; Meng, Zhang, and Ji 2023). We imaged two planes, a superficial plane 1 (P1, 80-120µm from pia) and a deep plane 2 (P2), with a difference of 100 µm between the center of each plane (Fig. 5F). We found that the mean baseline PW-evoked response strength for VIP interneurons was significantly larger than the low & moderate responding L2/3 neuronal population but was significantly lower than the high responders (Fig. 5B, VIP=0.11±0.009, n=302 cells, L2/3 neurons=0.03±0.001, n=1058 cells, high responders=0.87±0.13, n=41 cells, Two-way ANOVA, P<0.0001). We did not observe a significant change in the mean response strength when comparing VIP interneurons pre- and post-PRWS (Fig. 5C, pre=PRWS=0.11±0.009, post-PRWS=0.11±0.008, n=302 cells, Paired t test, P=0.7).

**Figure 5:**
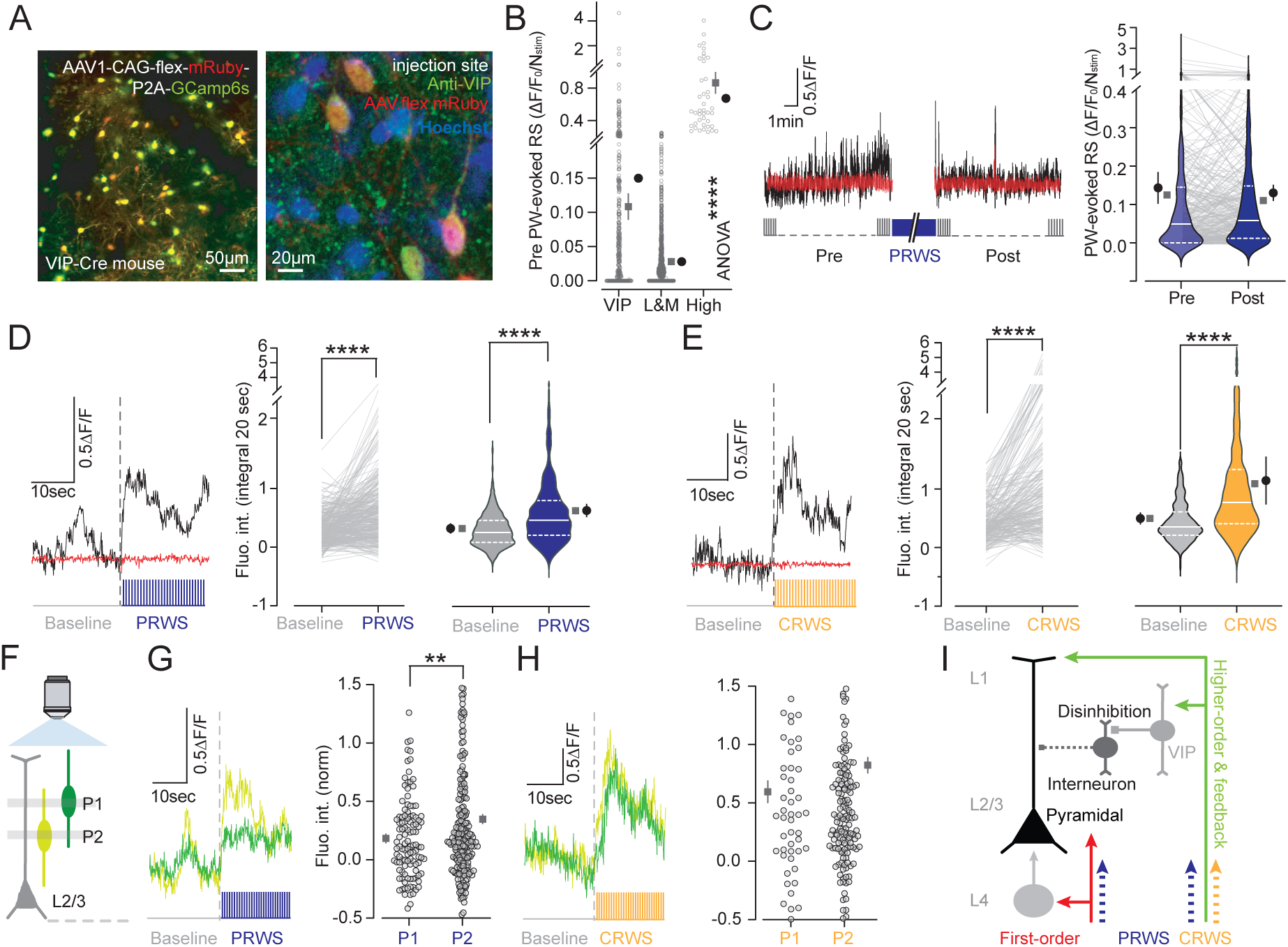
RWS non-selectively activates VIP interneurons. **(A) Left,** Example 2PLSM image of flex.mRuby.GCaMP6s-expressing VIP interneurons in the VIP-Cre mouse line. **Right,** representative confocal image after post-hoc anti-VIP immunocytochemistry on slices of barrel cortex from 2PLSM imaged VIP-Cre mice (green, anti-VIP; red, AAV1.CAG.Flex.mRuby.P2A.GCaMP6s; blue, Hoechst staining). **(B)** Pre-RWS PW-evoked response strength (RS) of VIP interneurons (n=341 cells, N=7 mice), and low & moderate (n=1058 cells, N=11 mice) and high (n=41, N=11 mice, one-way ANOVA, ****P<0.0001) responding L2/3 neurons. Squares and circles represent the means ± SEM over cells and mice, respectively. **(C) Left,** example trace of GCaMP6s (black) or mRuby fluorescence from a VIP interneuron, pre- and post-PRWS. **Right,** pre- and post-PRWS PW-evoked RS of VIP neurons (Paired t test, n=341 cells, P=0.2; N=7 mice, P=0.45). Grey lines, paired responses. Violin plots depict median (solid) and quartiles (dotted) bars. **(D, E), Left,** example trace of GCaMP6s fluorescence (black) or mRuby (integrated over 20s) from a VIP interneuron before and during PRWS **(D)** or CRWS **(E)**. **Right**, paired response and violin plots of normalized fluorescence intensity (norm.) during baseline, PRWS (Paired t-test, n=341 cells, ****P<0.0001; N=7 mice, P=0.047) or CRWS (Paired t-test, n=231 cells, ****P<0.0001; N=5 mice, P=0.08). **(F)** VIP interneurons were imaged at two planes in upper layers of S1, plane (P) 1 is closest to the pia and P2 is 100µm below. **(G, H) Left,** average VIP interneuron GCaMP6s fluorescence for P1 (darker green) and P2 (light green) for baseline (integrated over 20s) and during PRWS **(G)** or CRWS **(H)**. **Right,** normalized integrated fluorescence intensities duirng PRWS **(G**, P1 vs P2, Paired t test, **P=0.006) or CRWS **(H,** P1 vs P2, Paired t test, P=0.18). **(I)** Circuit diagram summarizing the RWS-evoked plasticity model. PRWS (blue) activates first-order thalamocortical (TC; red) as well as higher-order TC and feedback inputs (green), which activate disinhibitory VIP interneurons (grey). These combined inputs drive a potentiation of PW-evoked responses and a recruitment of neuronal responsivity (28%). CRWS (orange) may only activate higher-order TC and feedback inputs, also activating disinhibitory VIP interneurons, but this is not sufficient to drive potentiation and favors suppression of neurons (30%).

To test if VIP interneurons were activated during the repetitive sensory stimulation, we measured their responses during both experimental conditions. We found that these cells exhibited a significant increase in activity for at least 20s during PRWS compared to a 20s-baseline period just prior (Fig. 5D, pre-PRWS=0.3±0.15, post-PRWS=0.6±0.03, n=341 cells, Paired t test, P<0.0001). Surprisingly, we also found a significant increase in VIP interneuron activity upon CRWS, which suggests that the activation of VIP interneurons may not be whisker-selective (Fig. 5E, pre-CRWS=0.4±0.02, and post-CRWS=1.07±0.07, n=231 cells, Paired t test P<0.001). Although this effect was on trend, it was not significant when analyzed over mice (n=5 mice, P=0.08, Table 1-1).

Comparing VIP interneurons at different depths (P1 and P2), we found that PRWS significantly activated deeper as compared to superficial VIP interneurons (Fig 5F, P1=0.18±0.38, n=117, P2=0.35±0.04, n=224 cells, Unpaired t test, P=0.006). In contrast, CRWS did not have the same effect (Fig. 5G, P1=0.52±0.92, n=53 cells; P2=0.69±0.05, n=178 cells, Unpaired t test, P=0.18). Overall, we found that PRWS or CRWS strongly activated VIP interneurons for a sustained period (>1 min), with PRWS preferentially activating deeper VIP interneurons as compared to CRWS. Initially, VIP interneurons were more responsive to PW stimulation than the overall L2/3 neuronal population, yet they are less responsive than the high responding L2/3 neurons. Nonetheless, VIP interneurons are not potentiated post RWS, whereas are large part of the L2/3 neurons’ activity is potentiated.

## DISCUSSION

Repeated whisker stimulation in rodents under anesthesia potentiates local field potentials and elicits long-term potentiation (LTP) of cortical excitatory synapses (Mégevand et al. 2009; Gambino et al. 2014; Han et al. 2015; Williams and Holtmaat 2019). Here, we rhythmically stimulated a single whisker in awake mice and demonstrated that this modulates sensory-evoked population activity in L2/3 of the somatosensory cortex, as measured by somatic Ca^2+^ signals, which correlates with spiking rates (Zhang et al. 2023).

We found that a bout of repetitive whisker stimulation induces a prolonged (1h) increase in subsequent whisker-evoked activity in L2/3 neurons that initially responded at low and moderate levels to whisker deflections (Fig. 1-3). This effect was selective for neurons in the home cortical column of the stimulated whisker (PRWS) and did not occur when a far-away surround whisker was stimulated (CRWS). This selectivity rules out that it was attributable to sensory stimulation protocol or the repeated imaging perse. In a separate experiment, we also determined that the overall whisking rates or stimulus-evoked whisking rates do not change upon the RWS protocol (extended data Fig. 1-1). Altogether, this strongly suggests that the selective modulation of activity upon PRWS is due to changes in the synaptic pathways that are associated with the principal whisker. The effects bear similarities to electrically and sensory-evoked LTP *in vivo* and in experience-dependent plasticity paradigms albeit we have not tested whether they are the direct result of the synaptic LTP mechanisms that were previously observed under anesthesia (Gambino et al. 2014; Glazewski et al. 1996; Margolis et al. 2012; Han et al. 2015; Williams and Holtmaat 2019).

PRWS recruited more cells to the whisker-responding pool, whereas CRWS led to relatively more cells with suppressed activity. Nonetheless, PRWS also induced a decrease in activity rates in a subgroup of neurons, even to the extent that some stopped being responsive (suppressed) to whisker deflections altogether. A small population of L2/3 neurons that initially responded robustly decreased their activity, irrespective of PRWS or CRWS, which is a phenomenon that has also been observed in experience-dependent plasticity paradigms (Margolis et al. 2012). Together, these data indicate that the neuronal population has a bidirectional sensitivity to the repeated sensory stimulation. In contrast to the neurons that displayed moderate baseline activity, those with either very high or very low response rates were prone to lower their activity or lose it, respectively. This decrease in activity may be the result of synaptic weakening or homeostatic plasticity, possibly induced by asynchronous pre and postsynaptic activity in some neurons that can cause synaptic depression, or by increased levels of inhibition that normalize neuronal activity, respectively (Jacob et al. 2007; Knott et al. 2002). Conversely, the potentiation of the sensory-evoked activity could be aided by disinhibitory mechanisms, which are also known to support synaptic plasticity in the barrel cortex (Gambino and Holtmaat 2012; Williams and Holtmaat 2019; Li et al. 2014; Letzkus, Wolff, and Lüthi 2015). Differential effects of these opposing mechanisms may underlie the different fractions of neurons with persistent and suppressed responses as observed upon PRWS and CRWS. For example, for neurons with a low baseline response strength, the PRWS-evoked potentiation may counteract depression or homeostatic inhibition and protect them from being suppressed, whereas the CRWS, which does not potentiate responses, may fail in keeping these neurons in the responsive population.

We found that the potentiation lasts for ∼1hr on average which is relatively brief compared to long-term experience-dependent forms of plasticity in the sensory cortex. For example, whisker deprivation, which causes a sustained change in input to the sensory cortex has been shown to result in functional and even structural plasticity lasting over days to weeks (Margolis et al. 2012; Wilbrecht et al. 2010). Similarly, a period of environmental enrichment occludes subsequent RWS-induced plasticity (Mégevand et al. 2009). Nonetheless, the short-term plasticity that we observed might represent initial plasticity effects of an abrupt change in sensory experience that could be reinforced if it were continued, leading to long-term changes in the synaptic circuit.

PRWS could potentiate the activity of neurons even when their baseline whisker-evoked activity levels were low or absent (Fig. 1), suggesting that responsivity and plasticity are poorly correlated. Indeed, when comparing the level of RWS-evoked activity with the increase in whisker-evoked responses post-RWS, we did not observe a significant correlation (Fig. 3, 4). Instead, we found an inverse correlation between the initial whisker-evoked response strength and the level of activity during RWS. This shows that the level of plasticity could be independent of spike rates and that RWS can evoke plasticity in neurons that are normally poorly responsive to whisker deflections. These observations are in line with previous findings in that synaptic LTP can be independent of spiking and rely on sustained sub-threshold depolarizations, which may not be reflected in the somatic Ca^2+^ signals detected here (Golding, Staff, and Spruston 2002; Gambino et al. 2014; Lavzin et al. 2012; Brandalise et al. 2022).

In contrast to L2/3 neurons, VIP interneurons are highly sensitive to active whisking and touch (Yu et al. 2019; Lee et al. 2013), which we also observed during RWS (Fig. 5). VIP interneurons form well-characterized disinhibitory motifs for pyramidal neuron dendrites through inhibition of SST interneurons and may facilitate sensory-evoked evoked LTP (Lee et al. 2013; Pfeffer et al. 2013; Pi et al. 2013; Williams and Holtmaat 2019). The local disinhibition of dendrites during RWS could facilitate plasticity independent of parent neuronal activity levels, which provides another mechanism for the absence of a correlation between RWS-evoked activity and plasticity. Interestingly, we found that in contrast to the L2/3 neurons, the increase in RWS-evoked VIP interneuron activity was not whisker-selective and did not lead to a potentiation of subsequent single whisker-evoked VIP interneuron responses (Fig. 5), which corroborates our interpretation that the potentiation of L2/3 activity was due to changes in L2/3-associated synaptic pathways, and not attributable to post-RWS alterations in active whisking or active touch.

VIP interneuron subtypes are distributed over the cortex, hence superficial and deeper cells may on average have different morphological and functional signatures and may be implicated in different synaptic connectivity motifs (Prönneke et al. 2020; Jiang et al. 2015; Gouwens et al. 2020). To date, intralayer differences in VIP interneuron activity during sensory stimulation has not been described. Here, extended depth of field imaging has allowed us to examine activity in the superficial (P1, 80-120µm from pia) versus a deep (P2) plane 100µm below respectively (Fig 5). We found that the somewhat deeper VIP interneurons were more strongly activated than the very superficial neurons. This deeper pool of VIP interneurons could contain the L2 bipolar interneurons that constitute disinhibitory motifs (Prönneke et al. 2020; Georgiou et al. 2022). However, to make more precise inferences, a detailed morphological, functional or connectivity analysis would have to be performed. Nonetheless, the data do suggest that VIP interneurons are broadly activated and may cause disinhibition over a wide area of the somatosensory cortex, which is congruent with the fact that non-selective higher-order thalamocortical, cholinergic, and cortico-cortical circuits activate them (Fig. 5I) (Lee et al. 2013; Pi et al. 2013; Williams and Holtmaat 2019; Fu et al. 2014; Yu et al. 2019; Gambino et al. 2014).

Altogether, our findings paint the following picture: RWS increases activity rates of L2/3 neurons and increases the responsivity of individual neurons to subsequent sensory stimulation in a whisker-selective manner. VIP interneurons only increase their activity during the RWS stimulation period in a whisker non-selective manner. The large-scale disinhibition caused by the activation of VIP interneurons during RWS could open a gate for the potentiation of whisker-selective synaptic circuits on L2/3 neurons. Since VIP interneurons themselves were not potentiated, this excludes the possibility that protracted VIP-mediated disinhibition was directly responsible for the observed increase in L2/3 neuronal activity. Considering the synaptic circuits, we speculate that PRWS activates both first-order & higher-order thalamocortical (TC) and feedback inputs (Fig. 5l). These combined inputs may activate disinhibitory VIP interneurons and drive the potentiation of PW-evoked responses. CRWS, on the other hand, may only activate higher-order TC and feedback inputs, as well as disinhibitory VIP interneurons. However, this is not sufficient to drive potentiation without the activation of neurons through first-order TC inputs, and thus favors the suppression of neuronal responses.

Overall, this work indicates that the cortical representation of sensory input is dynamic and can be modulated over an extended period by repetitive sensory stimulation, via mechanisms that may involve activation of whisker-independent, barrel cortex-wide disinhibitory circuit motifs. In future endeavors, it will be important to test how modulating VIP interneuron activity shapes these population dynamics. It will also be interesting to determine if such sensory-driven mechanisms of plasticity also underly the reshaping of cortical representations and receptive fields during sensory deprivation-mediated plasticity, in disease, or during the formation of neuronal ensembles upon sensory learning (Hamdy et al. 1998; Kowalewski et al. 2012; Rose et al. 2016).

## Acknowledgements

We thank Ronan Chéreau, Céline Dürst, and Foivos Markopoulos for their technical assistance and critical evaluation of the experimental design; and Ronan Chéreau for help with software and comments on the manuscript. This work was supported by the Swiss National Science Foundation (grants # 31003A_153448 and 31003A_173125, 31003A_204562 to A.H.), and the International Foundation for Research in paraplegia (A.H.), and the Brain and Behavior Research Foundation (NARSAD YIG #28782 to L.E.W), and the Biotechnology and Biological Sciences Research Council (BB/V005405/1 to L.E.W).

---

**Figure 1-1:** Whisker movements during the stimulus protocol. **(A)** Calculating the whisker movement index (MI, arbitrary units, a.u.). Whiskers ipsi- (purple) and contralateral (pink) to the capillary tube were imaged at 112 Hz using a CCD digital camera placed under the snout of mice. To extract whisker movement, ROIs were drawn, from which the whisker position of each individual frame (orange) was correlated to the average whisker position across the entire movie (green). **(B)** Calculating mean overall whisker movement. Top, MI of the ipsi- and contralateral whiskers across the 10-minute protocol pre (red) and post RWS (blue) for 1 mouse. Stimulations are marked in grey. Bottom, normalized mean MI for the ipsi- and contralateral whiskers of 4 mice Pre- and Post-RWS. (ipsi pre=1.0, post=0.98±0.03; contra pre=1.19±0.11, post=1.12±0.09, one-way ANOVA, P=0.24). **(C)** Calculating stimulus-evoked whisker movement pre- and post-RWS. Top, schematic illustrating the calculation of the average MI 2 (s, 224 frames) before the start of a stimulus (from dashed box in B) and 2s after the end of the stimulus. Middle, normalized mean MI for each mouse for ipsi- (left, pre before=1.0, after=0.86±0.06; post before=0.93±0.01, after=0.90±0.07, one-way ANOVA, P=0.25) and contralateral (right, pre before=1.0, after=0.93±0.05; post before=0.95±0.05, after=0.89±0.06, one-way ANOVA, P=0.44) whiskers. Bottom left, scatterplot comparing the MI of ipsilateral whiskers before and after stimulus presentation pre- (n=236 stims, r=0.47, P<0.0001) and post-RWS (n=236 stims, r=0.47, P<0.0001). Bottom right, scatterplot comparing the MI of contralateral whiskers before and after stimulus presentation pre- (n=236 stims, r=0.43, P<0.0001) and post-RWS (n=236 stims, r = 0.44, P<0.0001).

