## Extended data figure 1-1 for "Repetitive sensory stimulation potentiates and recruits sensory-evoked cortical population activity"

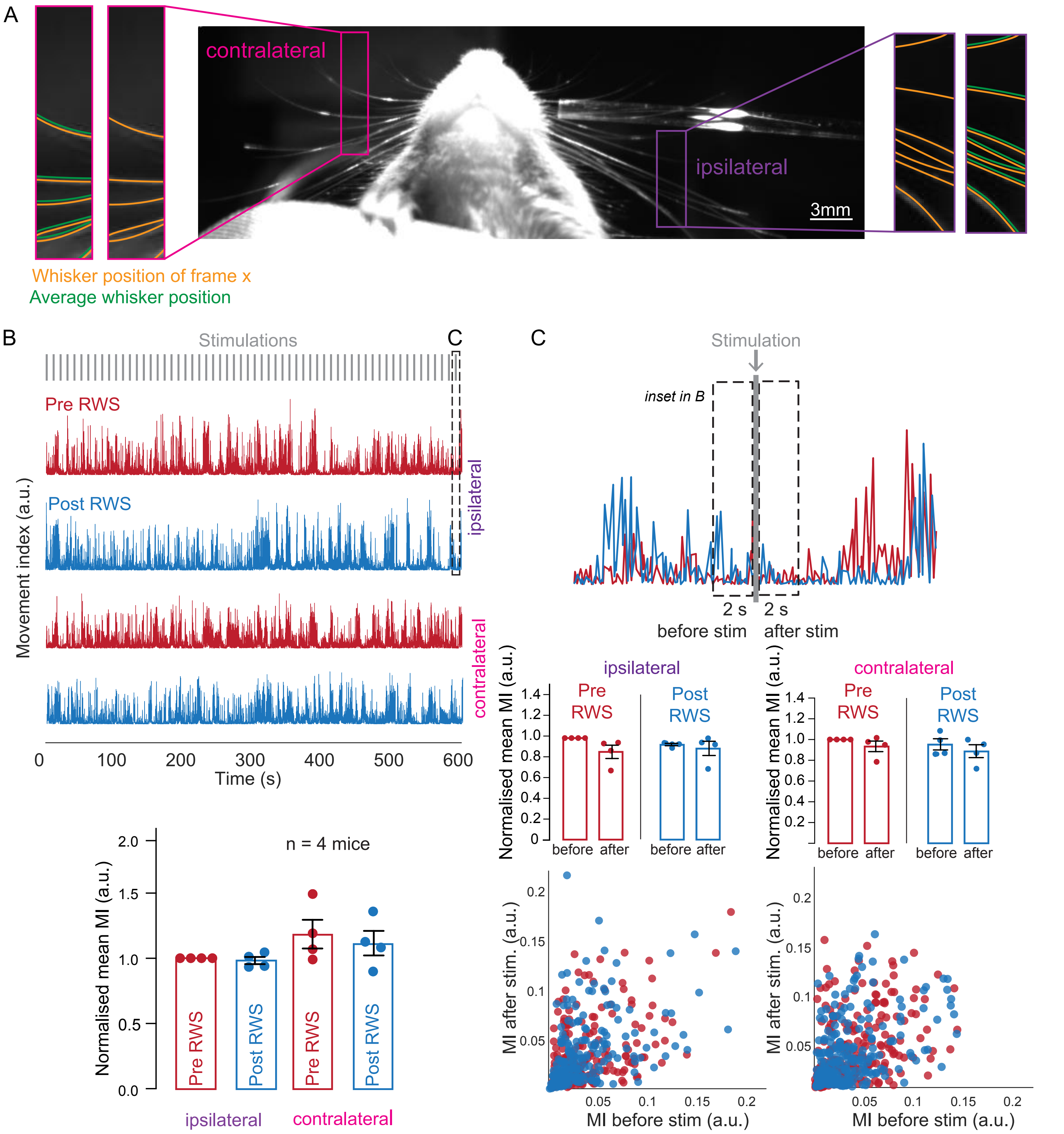


**Figure 1-1: Whisker movements during the stimulus protocol. (A)** Calculating the whisker movement index (MI, arbitrary units, a.u.). Whiskers ipsi- (purple) and contralateral (pink) to the capillary tube were imaged at 112 Hz using a CCD digital camera placed under the snout of mice. To extract whisker movement, ROIs were drawn, from which the whisker position of each individual frame (orange) was correlated to the average whisker position across the entire movie (green). **(B)** Calculating mean overall whisker movement. Top, MI of the ipsi- and contralateral whiskers across the 10-minute protocol pre (red) and post RWS (blue) for 1 mouse. Stimulations are marked in grey. Bottom, normalised mean MI for the ipsi- and contralateral whiskers of 4 mice Pre- and Post-RWS. (ipsi pre=1.0, post=0.98±0.03; contra pre=1.19±0.11, post=1.12±0.09, one-way ANOVA, P=0.24). **(C)** Calculating stimulus-evoked whisker movement pre- and post-RWS. Top, schematic illustrating the calculation of the average MI 2s (224 frames) before the start of a stimulus (from dashed box in B) and 2s after the end of the stimulus. Middle, normalised mean MI for each mouse for ipsi- (left, pre before=1.0, after=0.86±0.06; post before=0.93±0.01, after=0.90±0.07, one-way ANOVA, P=0.25) and contralateral (right, pre before=1.0, after=0.93±0.05; post before=0.95±0.05, after=0.89±0.06, one-way ANOVA, P=0.44) whiskers. Bottom left, scatterplot comparing the MI of ipsilateral whiskers before and after stimulus presentation pre- (n=236 stims, r=0.47, P<0.0001) and post-RWS (n=236 stims, r=0.47, P<0.0001). Bottom right, scatterplot comparing the MI of contralateral whiskers before and after stimulus presentation pre- (n=236 stims, r=0.43, P<0.0001) and post-RWS (n=236 stims, r = 0.44, P<0.0001).
