## Extended data table 1-1 for "Repetitive sensory stimulation potentiates and recruits sensory-evoked cortical population activity"

### Table 1-1. Descriptive Statistics

| Panel | test | n=cells, N=mice | *P* value | Other/units | mean 1 | Error (±SEM) | mean 2 | Error (SEM) | conclusion |
|---|---|---|---|---|---|---|---|---|---|
| Fig. 1E | Paired t test | 1099 cells | 0.0015 | $(\Delta F/F_0)$/Nstim | 0.06 | 0.007 | 0.07 | 0.005 | ** |
| | | 11 mice | 0.51 | | 0.06 | 0.01 | 0.07 | 0.009 | ns |
| Fig. 1F | Paired t test | 829 cells | 0.4 | $(\Delta F/F_0)$/Nstim | 0.076 | 0.008 | 0.072 | 0.008 | ns |
| | | 11 mice | 0.58 | | 0.076 | 0.01 | 0.08 | 0.016 | ns |
| Fig. 1G | Simple linear regression | 11 mice, PRWS=1099 cells CRWS=829 cells | P<0.0001 | | slope=0.62 | | slope=0.85 | | **** |
| Fig. 1H | Iterative Grubb's outlier test, α=0.0001 | 11 mice, PRWS=41 cells CRWS=37 cells | | $(\Delta F/F_0)$/Nstim | 0.87 | 0.13 | 0.72 | 0.06 | outliers |
| Fig. 1I left | Paired t test | 1058 cells | P<0.0001 | $(\Delta F/F_0)$/Nstim | 0.03 | 0.001 | 0.05 | 0.003 | **** |
| | | 11 mice | 0.006 | | 0.029 | 0.002 | 0.05 | 0.006 | ** |
| Fig. 1I right | Paired t test | 792 cells | 0.26 | $(\Delta F/F_0)$/Nstim | 0.04 | 0.002 | 0.045 | 0.004 | ns |
| | | 11 mice | 0.67 | | 0.045 | 0.005 | 0.047 | 0.006 | ns |
| Fig. 1J right | Paired t test | 41 cells | 0.0008 | $(\Delta F/F_0)$/Nstim | 0.87 | 0.13 | 0.60 | 0.09 | *** |
| | | 11 mice | 0.04 | | 0.75 | 0.11 | 0.53 | 0.10 | * |
| Fig. 1J left | Paired t test | 37 cells | <0.0001 | $(\Delta F/F_0)$/Nstim | 0.84 | 0.11 | 0.66 | 0.12 | **** |
| | | 11 mice | 0.0001 | | 0.76 | 0.14 | 0.55 | 0.14 | *** |
| Fig. 1 K | Mixed-effects model | 11 mice | 0.0004 | Pre/Post (%) | F(1,39)=15.1 | | | | *** |
| | Uncorrected Fisher's LSD | | 0.0004 | | PRWS Low/mod= 190.9% | 37.85 | PRWS Hi= 74.65% | 12.94 | *** |
| | | | 0.10 | | CRWS Low/mod= 116.3% | 11.88 | CRWS Hi= 64.52% | 6.24 | ns |
| | | | 0.018 | | PRWS Low/mod = 190.9% | 37.85 | CRWS Low/mod = 116.3% | 11.88 | * |
| | | | 0.74 | | Hi PRWS= 74.65% | 12.94 | Hi CRWS= 64.52% | 6.24 | ns |
| Fig. 2 A middle | Paired t test | 465 cells | <0.0001 | $(\Delta F/F_0)$/Nstim | 0.05 | 0.002 | 0.08 | 0.006 | **** |
| | | 11 mice | 0.0003 | | 0.06 | 0.004 | 0.09 | 0.008 | *** |
| Fig. 2 A right | Paired t test | 279 cells | 0.06 | $(\Delta F/F_0)$/Nstim | 0.076 | 0.005 | 0.09 | 0.009 | ns |
| | | 11 mice | 0.15 | | 0.08 | 0.006 | 0.096 | 0.008 | |
| Fig. 2 B | Unpaired t test | PRWS= 307 cells CRWS=131 cells | 0.14 | $(\Delta F/F_0)$/Nstim | 0.05 | 0.004 | 0.07 | 0.001 | ns |
| | | 11 mice | 0.53 | | 0.04 | 0.007 | 0.05 | 0.01 | |
| Fig. 2 C | Unpaired t test | PRWS= 205 cells CRWS=258 cells | <0.0001 | $(\Delta F/F_0)$/Nstim | 0.03 | 0.002 | 0.04 | 0.003 | **** |
| | | 11 mice | 0.037 | | 0.025 | 0.004 | 0.033 | 0.005 | * |
| Fig. 2 D | Chi square test, Chi-square=19.8 , DF=4 | 11 mice, PRWS=1099 cells CRWS=829 cells | 0.0005 | persistent | 42% (n=465) | | 34% (n=279) | | *** |
| | | | | recruited | 28% (n=307) | | 16% (n=131) | | |
| | | | | suppressed | 19% (n=205) | | 31% (n=258) | | |
| | | | | no response | 7% (n=81) | | 15% (n=124) | | |
| | | | | hi responders | 4% (n=41) | | 4% (n=37) | | |
| Fig. 3 B | Two-way RM ANOVA | PRWS= 382 cells CRWS=304 cells | 0.0006 | $(\Delta F/F_0)$/Nstim | | | | | *** |
| | Two-way RM ANOVA | 6 mice | 0.026 | | | | | | * |
| Fig. 3 B | Dunnett's multiple comparisons | PRWS= 382 cells | <0.0001 | $(\Delta F/F_0)$/Nstim | -10 min=0.03 | 0.002 | 10min= 0.045 | 0.005 | **** |
| | | | 0.0001 | | | | 60min=0.040 | 0.040 | *** |
| | | | 0.51 | | | | 120min=0.032 | 0.003 | ns |
| | | | 0.40 | | | | 180min=0.034 | 0.004 | ns |
| | Uncorrected Fisher's LSD | 6 mice | 0.005 | | -10 min=0.029 | 0.003 | 10min= 0.046 | 0.008 | ** |
| | | | 0.059 | | | | 60min=0.040 | 0.007 | ns |
| | | | 0.56 | | | | 120min=0.032 | 0.004 | |
| | | | 0.41 | | | | 180min=0.033 | 0.006 | |
| Fig. 3 B | Dunnett's multiple comparisons | CRWS= 304 cells | 0.9 | $(\Delta F/F_0)$/Nstim | -10 min=0.034 | 0.004 | 10min=0.034 | 0.005 | ns |
| | | | 0.3 | | | | 60min=0.040 | 0.005 | |
| | | | 0.7 | | | | 120min=0.033 | 0.004 | |
| | | | 0.5 | | | | 180min=0.032 | 0.004 | |
| | | 6 mice | 0.6 | | -10 min=0.035 | 0.004 | 0.038 | 0.006 | ns |
| | | | 0.3 | | | | 0.029 | 0.005 | |

| | | | | | | | | | |
|---|---|---|---|---|---|---|---|---|---|
| | Uncorrected Fisher's LSD | | 0.5 | | | | 0.031 | 0.005 | |
| | | | 0.4 | | | | 0.030 | 0.004 | |
| Fig. 3 C | Two-way RM ANOVA | PRWS= 382 cells CRWS=304 cells | <0.0001 | #events/Nstim | | | | | **** |
| | | N=6 mice | 0.026 | | | | | | * |
| Fig. 3 C | Dunnett's multiple comparisons | PRWS= 382 cells | <0.0001 | #events/Nstim | -10 min=0.04 | 0.003 | 10min=0.056 | 0.005 | **** |
| | | | 0.001 | | | | 60min=0.054 | 0.001 | ** |
| | | | 0.7 | | | | 120min=0.04 | 0.004 | ns |
| | | | 0.99 | | | | 180min=0.04 | 0.004 | ns |
| | Uncorrected Fisher's LSD | 6 mice | 0.013 | #events/Nstim | 0.049 | 0.004 | 0.065 | 0.007 | * |
| | | | 0.05 | | | | 0.063 | 0.006 | ns |
| | | | 0.77 | | | | 0.047 | 0.007 | ns |
| | | | 0.94 | | | | 0.049 | 0.005 | ns |
| Fig. 3 C | Dunnett's multiple comparisons | CRWS= 304 cells | 0.99 | #events/Nstim | -10 min=0.045 | 0.005 | 10min=0.046 | 0.006 | ns |
| | | | 0.3 | | | | 60min=0.040 | 0.005 | |
| | | | 0.2 | | | | 120min=0.040 | 0.005 | |
| | | | 0.3 | | | | 180min=0.040 | 0.005 | |
| | Uncorrected Fisher's LSD | 6 mice | 0.69 | #events/Nstim | 0.042 | 0.004 | 0.045 | 0.008 | ns |
| | | | 0.13 | | | | 0.037 | 0.004 | |
| | | | 0.06 | | | | 0.036 | 0.006 | |
| | | | 0.09 | | | | 0.037 | 0.005 | |
| Fig. 3 D | Two-way RM ANOVA | PRWS= 382 cells CRWS=304 cells | <0.0001 | $\Delta F/F_0$ | | | | | **** |
| | | 6 mice | 0.25 | | | | | | ns |
| Fig. 3 D | Dunnett's multiple comparisons | PRWS= 382 cells | 0.01 | $\Delta F/F_0$ | -10 min=0.04 | 0.02 | 10min=0.48 | 0.02 | * |
| | | | 0.002 | | | | 60min=0.5 | 0.02 | ** |
| | | | 0.8 | | | | 120min=0.040 | 0.003 | ns |
| | | | 0.4 | | | | 180min=0.44 | 0.02 | ns |
| | Uncorrected Fisher's LSD | 6 mice | 0.09 | | 0.41 | 0.05 | 0.47 | 0.07 | ns |
| | | | 0.006 | | | | 0.51 | 0.05 | ** |
| | | | 0.77 | | | | 0.44 | 0.04 | ns |
| | | | 0.24 | | | | 0.46 | 0.04 | ns |
| Fig. 3 D | Dunnett's multiple comparisons | CRWS= 304 cells | 0.2 | $\Delta F/F_0$ | -10 min=0.04 | 0.03 | 10min=0.36 | 0.02 | ns |
| | | | 0.6 | | | | 60min=0.37 | 0.03 | ns |
| | | | 0.1 | | | | 120min=0.47 | 0.03 | ns |
| | | | 0.5 | | | | 180min=0.45 | 0.03 | ns |
| | Uncorrected Fisher's LSD | 6 mice | 0.76 | | 0.37 | 0.07 | 0.36 | 0.07 | ns |
| | | | 0.59 | | | | 0.35 | 0.06 | ns |
| | | | 0.01 | | | | 0.44 | 0.05 | * |
| | | | 0.05 | | | | 0.42 | 0.05 | ns |
| Fig 3E | Two-way RM ANOVA | PRWS 162 cells | 0.36 | $(\Delta F/F_0)$/Nstim | -10 min=0.05 | 0.02 | 24 hrs=0.06 | 0.01 | ns |
| | | CRWS 134 cells | | | 0.04 | 0.006 | 0.06 | 0.01 | |
| | | 3 mice | 0.054 | | -10 min=0.06 | 0.02 | 0.06 | 0.02 | |
| | | 2 mice | | | 0.05 | 0.005 | 0.05 | 0.001 | |
| Fig. 4 A | Paired t test | 260 cells | <0.0001 | $\Delta F/F_0$ integrated over 20 sec | baseline=0.2 | 0.02 | PRWS=0.3 | 0.02 | **** |
| | | 3 mice | 0.027 | | 0.18 | 0.13 | 0.30 | 0.11 | * |
| Fig. 4 B | Paired t test | 115 cells | 0.6 | $\Delta F/F_0$ integrated over 20 sec | baseline=0.47 | 0.02 | CRWS=0.5 | 0.03 | ns |
| | | 3 mice | 0.8 | | 0.47 | 0.03 | 0.5 | 0.08 | |
| Fig. 4 C | Pearson r correlation | 3 mice 115 cells | 1.0 | $\Delta F/F_0$ integrated over 20 sec vs. post/pre% | r=0.0002 | linear regressio n | slope=0.089 | nonzero slope? P=0.99 | ns |
| Fig. 4 D | Pearson r correlation | 3 mice 115 cells | 0.0001 | $\Delta F/F_0$ integrated over 20 sec vs. $(\Delta F/F_0)$/Nstim | r=-0.353 | simple linear regressio n | slope=-0.008 | nonzero slope? P=0.0001 | **** |
| Fig. 4 C | Pearson r correlation | 3 mice 115 cells | 1.0 | $\Delta F/F_0$ integrated over 20 sec vs. post/pre% | r=0.0002 | linear regressio n | slope=0.089 | nonzero slope? P=0.99 | ns |
| Fig. 4 C inset | Paired t test | 3 mice 115 cells | 0.027 | $(\Delta F/F_0)$/Nstim | pre=0.027 | 0.002 | post=0.031 | 0.002 | * |
| Fig. 4 D | Pearson r correlation | 3 mice 115 cells | 0.0001 | $\Delta F/F_0$ integrated over 20 sec vs. $(\Delta F/F_0)$/Nstim | r=-0.353 | simple linear regressio n | slope=-0.008 | nonzero slope? P=0.0001 | **** |
| Fig. 5 B | two-way ANOVA | 341 cells | <0.0001 | VIP | 0.11 | 0.009 | | | **** |
| | | 1058 cells | | Low/mod | 0.03 | 0.001 | | | |
| | | 41 cells | | Hi responders | 0.87 | 0.13 | | | |
| | two-way ANOVA | 7 mice | <0.0001 | VIP | 0.15 | 0.04 | | | |
| | | 11 mice | | Low/mod | 0.03 | 0.002 | | | |
| | | 11 mice | | Hi responders | 0.75 | 0.11 | | | |
| Fig. 5 C | Paired t test | 341 cells | 0.22 | $(\Delta F/F_0)$/Nstim | 0.13 | 0.017 | 0.119 | 0.011 | ns |
| | | 7 mice | 0.45 | | 0.14 | 0.04 | 0.13 | 0.02 | |
| Fig. 5 D | Paired t test | 341 cells | <0.0001 | $(\Delta F/F_0)$/Nstim | 0.30 | 0.015 | 0.596 | 0.028 | **** |
| | | 7 mice | 0.047 | | 0.31 | 0.07 | 0.61 | 0.099 | * |

| | | | | | | | | | |
|---|---|---|---|---|---|---|---|---|---|
| Fig. 5 E | Paired t test | 231 cells | <0.0001 | $\Delta F/F_0$ integrated over 20 sec | 0.43 | 0.022 | 1.065 | 1.065 | **** |
| | | 5 mice | 0.08 | | 0.04 | 0.09 | 1.15 | 0.38 | ns |
| Fig. 5G | Unpaired t test | 7 mice<br>P1=117 cells<br>P2=224 cells | 0.006 | Fluo intensity (norm to baseline) | P1= 0.18 | 0.038 | P2= 0.38 | 0.039 | ** |
| Fig. 5H | Unpaired t test | 5 mice<br>P1=53 cells<br>P2=178 cells | 0.175 | Fluo intensity (norm to baseline) | P1=0.52 | 0.093 | P2=0.699 | 0.066 | ns |
